# Anti-phage islands force their target phage to directly mediate island excision and spread

**DOI:** 10.1101/218164

**Authors:** Amelia C. McKitterick, Kimberley D. Seed

## Abstract

To defend against their adversaries, bacteria and phage engage in cycles of adaptation and counter-adaptation that shape their mutual evolution^1–3^. *Vibrio cholerae*, the causative agent of the diarrheal disease cholera, is antagonized by phages in the environment as well as in human hosts^4,5^. The lytic phage ICP1 has been recovered from cholera patient stool and water samples over at least 12 years in Bangladesh^6–8^ and is consequently considered a persistent predator of epidemic *V. cholerae* in this region. In previous work, we demonstrated that mobile genetic elements called <u>p</u>hage-inducible chromosomal island-like <u>e</u>lements (PLEs) protect *V. cholerae* from ICP1 infection^7,9^. PLEs initiate their anti-phage response by excising from the chromosome, however, the mechanism and molecular specificity underlying this response are not known. Here, we show that PLE 1 encodes a large serine recombinase, Int, that exploits an ICP1-specific protein, PexA, as a recombination directionality factor (RDF) to sense and excise in response to ICP1 infection. We validate the functionality and specificity of this unique recombination system, in which the recombinase and RDF are encoded in separate genomes. Additionally, we show that PexA is also hijacked to trigger excision in PLEs found in *V. cholerae* isolates recovered decades ago. Our results uncover an aspect of the molecular specificity underlying the longstanding conflict between a single predatory phage and *V. cholerae* PLE and contribute to our understanding of the molecular arms race that drives long-term evolution between combatting phage and their bacterial hosts in nature.

---

The five PLEs identified in *V. cholerae* isolates spanning a >60-year collection period exhibit a common, ICP1-dependent response, which initiates with the integrated PLE excising from the chromosome and circularizing^9^ (Fig. 1a). PLE excision facilitates mobilization as PLEs are transduced following ICP1 infection, permitting their spread to *V. cholerae* recipients^9^. Although the underlying mechanisms are not known, PLEs block the phage replication program in what appears to be an ICP1-specific manner, as other tested cholera phages do not stimulate PLE circularization and are not blocked by PLEs^9^. Further, protection against ICP1 is absolute— PLE^+^ cells produce no progeny phage. The transmission costs imposed by PLEs are a significant burden to ICP1 in nature, as some ICP1 isolates have acquired a CRISPR-Cas system to target PLEs, which allows ICP1 to persist in spite of PLE^7,8^. In addition to what has been documented regarding PLE circularization following ICP1 infection under laboratory conditions^7,9^, we note that ICP1-dependent PLE circularization can be detected in cholera patient stool (Fig. 1b), underscoring that *V. cholerae* PLE responds to ICP1 infection during disease in humans. Due to the apparent specificity and conservation of PLE circularization in response to ICP1, we set out to characterize the mechanism governing this response.

**Fig. 1.**
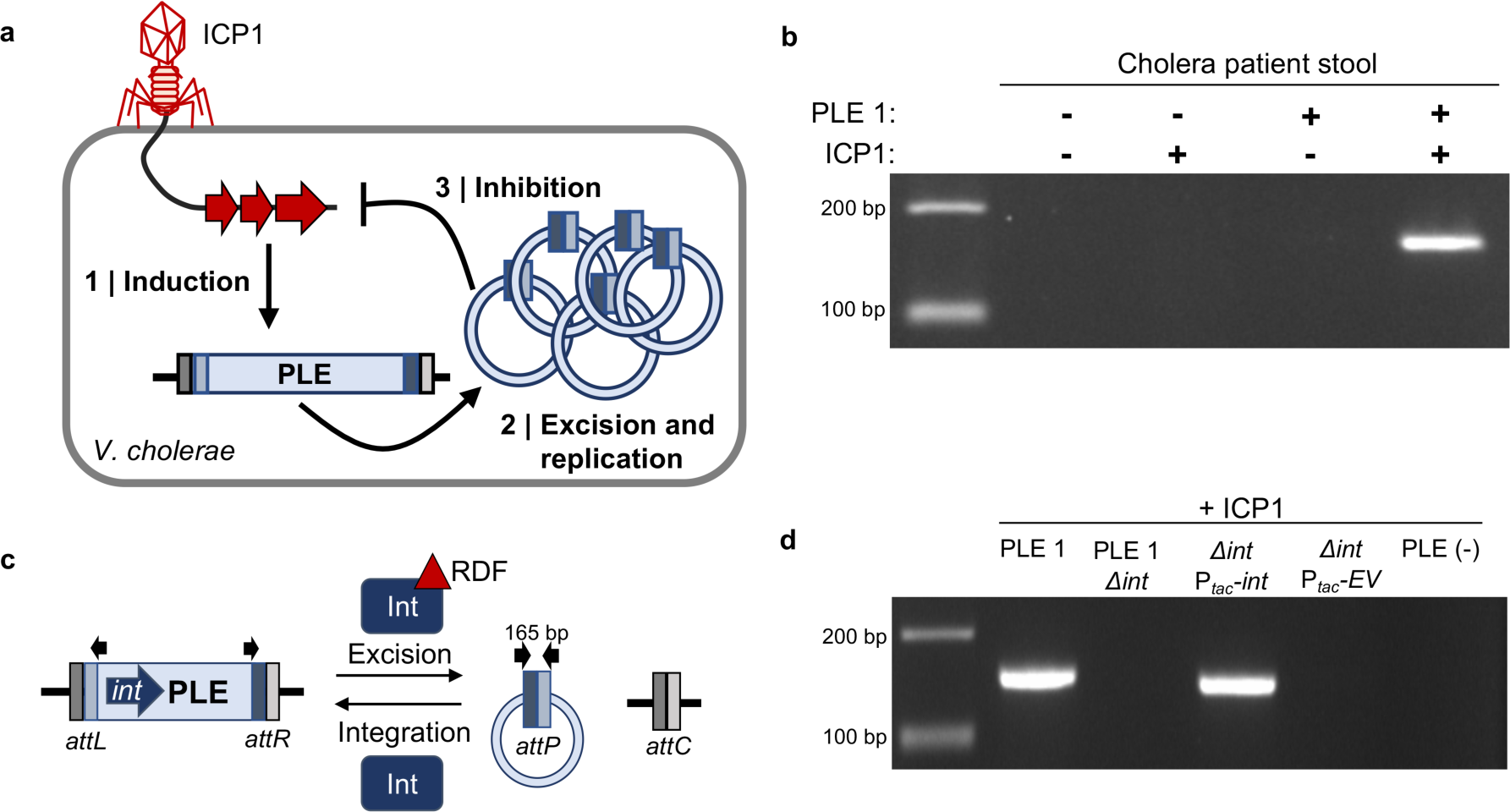
PLE circularizes during ICP1 infection and this response is dependent on PLE-encoded Int. **a**, ICP1 injects its DNA into a PLE(+) *V. cholerae* cell, leading to (1) PLE induction. (2) Induced PLE excises from the chromosome and circularizes, replicating to high copy number^9^. Through an unknown mechanism, (3) PLE inhibits the ICP1 replication program and protects the *V. cholerae* population from ICP1 predation. **b**, Circularized PLE, which is detected by PCR using outward-facing primers internal to the PLE, is found in cholera patient stool samples when PLE and ICP1 are both present. **c**, Model of LSR/RDF mediated integration and excision of PLE and the resulting *att* sites. Black arrows indicate primers used to detect PLE circularization. **d**, Detection of PLE 1 circularization during ICP1 infection. *V. cholerae* with the PLE 1 variant indicated was infected with ICP1 at a multiplicity of infection (MOI) of 5 and harvested 5m post infection to detect PLE circularization. Strains complemented with a plasmid harboring *int* under control of an IPTG inducible P_*tac*_ promoter or the empty vector (EV) control were induced 20m prior to phage infection. Uncropped gels are presented in Supplementary Fig. 5.

PLE encodes a gene product (Int) with an N-terminal serine recombinase domain and a large C-terminus containing a putative zinc ribbon domain and coiled-coiled motif characteristic of large serine recombinases (LSRs)^10^. Typically found in temperate phages, LSRs have the ability to catalyze recombination between attachment (*att*) sites^11–13^. Only the LSR is required to catalyze recombination between episomal (*attP*) and chromosomal (*attC*) sites, leading to integration of the episome into the host chromosome. To reverse this process, a recombination directionality factor (RDF) is required to physically interact with the LSR and direct the LSR to recombine the left (*attL*) and right (*attR*) attachment sites, resulting in excision of DNA between these sites (Fig. 1c). To determine if Int plays a role in PLE circularization during ICP1 infection, PLE 1 *Δint* was challenged with ICP1. Unlike wild-type PLE, circularization was not detected in the *Δint* strain. PLE circularization was restored with *in trans* Int expression, but only during ICP1 infection (Fig. 1d, Supplemental Fig. 1a). To determine if PLE 1 Int is a functional LSR, we performed integration assays to probe the ability of Int to recombine the *attC* and *attP* sites. Through both *in vivo* (Supplemental Fig. 1b) and *in vitro* (Supplemental Fig. 1c) assays, we found that Int is sufficient to recombine *attP* and *attC* sites, as is characteristic of LSRs^11,12^. These assays demonstrate that Int is necessary for PLE circularization in response to ICP1 infection and is a functional LSR that can catalyze recombination between *att* sites.

Expression of Int is not sufficient to catalyze circularization of PLE 1 in the absence of phage infection, so an RDF is required to direct Int to recombine the *attL* and *attR* sites as is characteristic of LSRs^14,15^. There are no conserved sequence characteristics of RDFs that enable homology-based identification^16^, however, in characterized LSR/RDF systems of temperate phages, both the LSR and RDF are encoded within the same genome^14,17^. To evaluate if the RDF is PLE-encoded, PLE 1 strains harboring gene cluster deletions of predicted open reading frames (ORFs) were screened for circularization defects during ICP1 infection. Unexpectedly, all of the PLE ORF knockouts still circularized, implying that the RDF is not PLE-encoded (Supplementary Fig. 2a). To establish the minimal PLE-encoded factors required for circularization, we constructed a ‘miniPLE’, which has Int under control of its endogenous promoter and a kanamycin cassette flanked by *att* sites, integrated into the *V. cholerae* chromosome in the same location as PLE 1 (Fig. 2a). In support of the mutational analyses (Supplemental Fig. 2a), the miniPLE circularized and excised from the chromosome during ICP1 infection (Fig 2a., Supplemental Fig. 2b). Together with the inability of PLE 1 *Δint* to circularize (Fig. 1d), these results demonstrate that Int is necessary and sufficient for PLE 1 circularization during ICP1 infection.

**Fig. 2.**
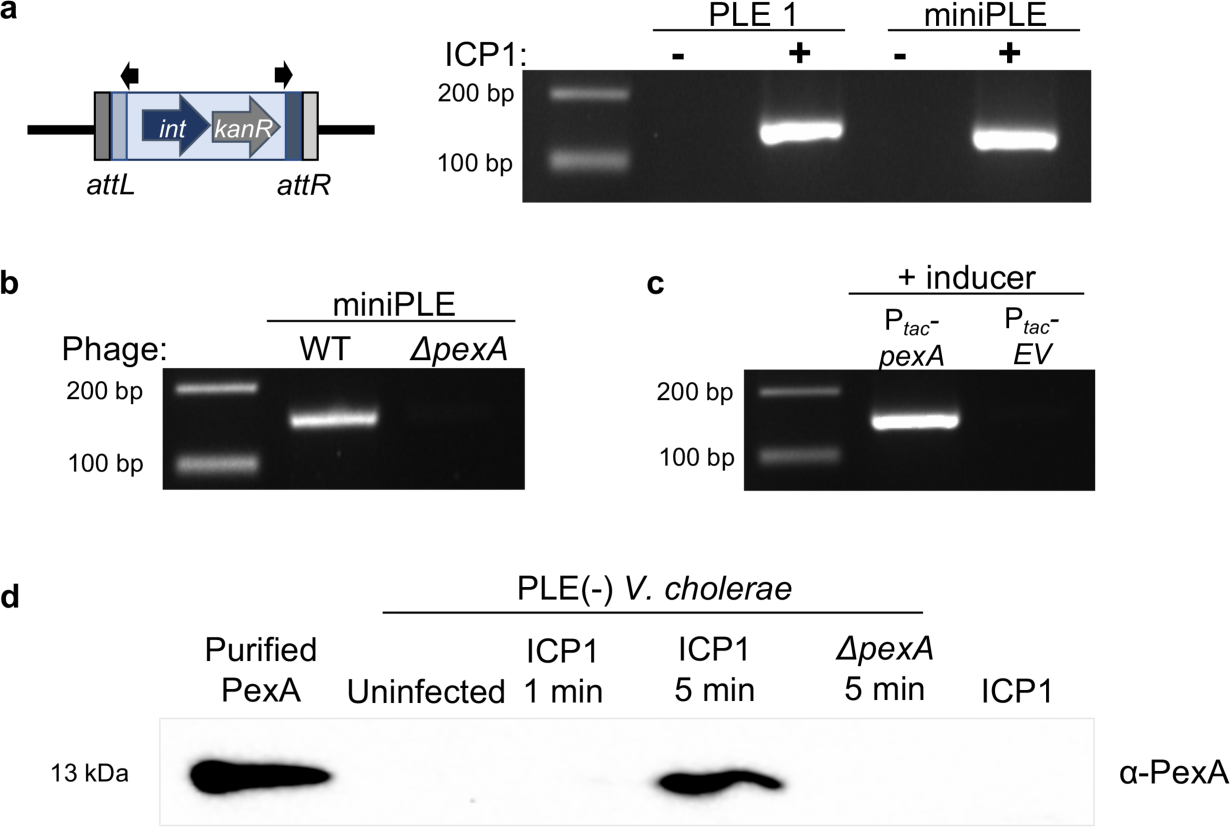
ICP1-encoded PexA is necessary and sufficient for PLE circularization. **a**, Cartoon of the miniPLE (left) containing PLE 1 *int* and a kanamycin cassette (*kanR*) integrated into the *V. cholerae* chromosome in the same location as PLE 1 with the same *att* sites. Circularization of miniPLE (right) can be detected 5m post ICP1 infection using the same primers as used for PLE 1 circularization. **b**, miniPLE circularization is not detected during ICP1 infection when *pexA* is knocked out. **c**, Circularization of miniPLE is detected in the absence of ICP1 infection when PexA is ectopically expressed. **d**, Western blot for PexA during ICP1 infection. PLE(-) *V. cholerae* cultures were probed for PexA at the listed times after infection with ICP1 or ICP1 *ΔpexA*. To determine if PexA is packaged in the phage particle, 5 times the PFU of phage as was used for infection was probed for the presence of PexA. Purified PexA (20 ng) was used as a positive control. Uncropped gels are presented in Supplementary Fig. 5.

Due to the specificity of circularization during ICP1 infection, we hypothesized that ICP1 encodes a gene product that directs Int-mediated PLE circularization during infection. To identify this phage-encoded gene product, we screened for ICP1 mutations that abolished miniPLE circularization during infection. Through this screen, we identified a hypothetical ICP1 gene product annotated as ORF51 (YP_004250992.1) that was dispensable for ICP1 plaque formation (Supplemental Fig. 3) but necessary for miniPLE circularization (Fig. 2b). Additionally, ectopic expression of this ICP1 gene product from a plasmid in the absence of phage infection resulted in robust miniPLE circularization (Fig. 2c). As this gene product is both necessary and sufficient for Int-mediated miniPLE excision, we named it <u>p</u>hage-<u>e</u>ncoded e<u>x</u>cisionase (PexA). PexA is a small protein unique to ICP1 isolates that has no sequence similarity to known proteins.

Consistent with the rapid kinetics of PLE 1 circularization^9^, we found that PexA is expressed *de novo* within five minutes of ICP1 infection (Fig. 2d), leading us to hypothesize that PexA is hijacked by PLE 1 to function as the RDF for Int-mediated PLE excision. The ability of PexA to physically interact with Int was probed with a bacterial adenylate cyclase two-hybrid (BACTH) assay, in which LacZ expression was detected when both Int and PexA were fused to adenylate cyclase subunits (Fig. 3a). This interaction was further validated using an *in vitro* pulldown assay, in which PexA coeluted with 6xHis-tagged Int (Fig. 3b).

**Fig. 3.**
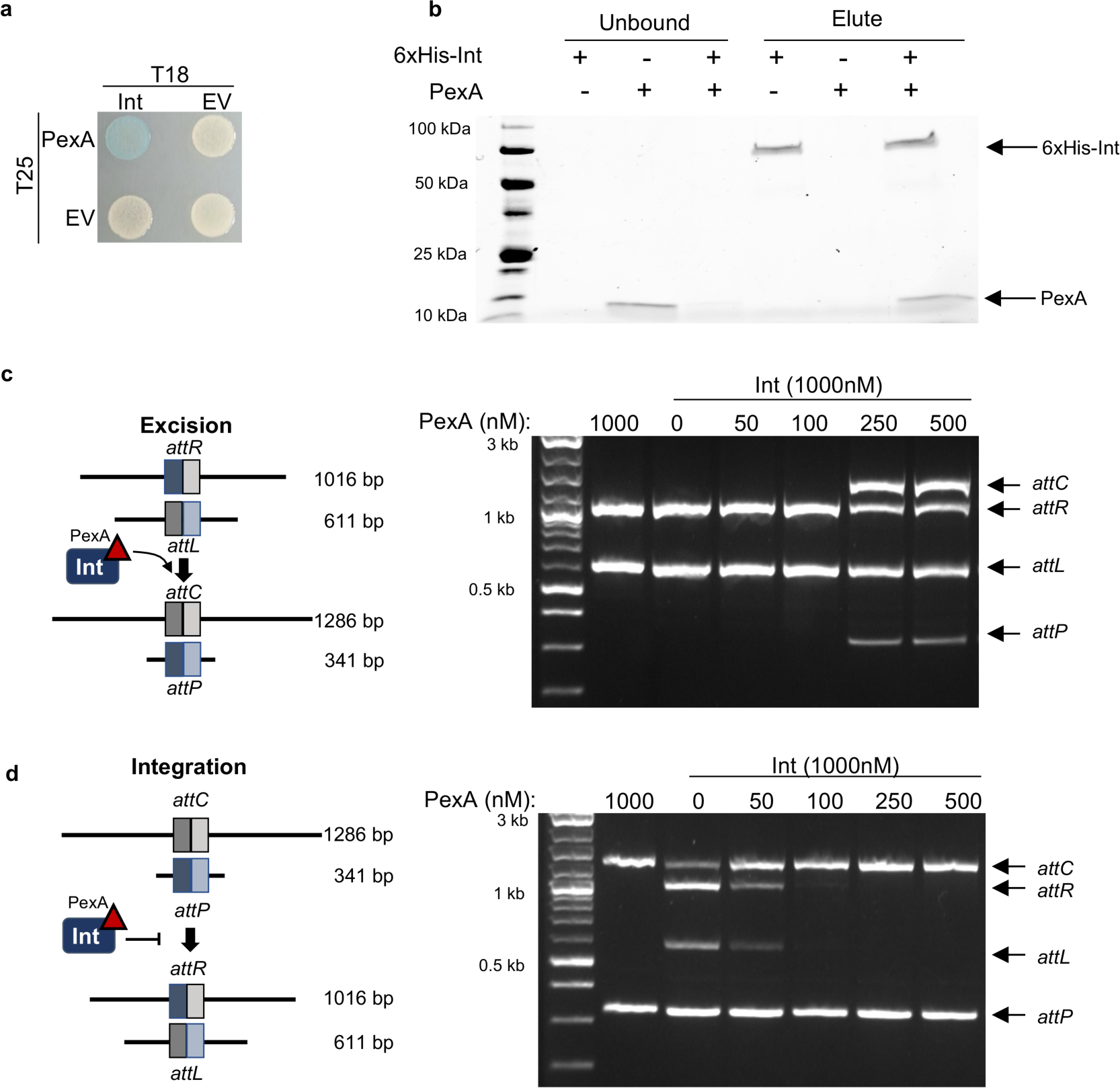
PLE exploits phage encoded PexA as the RDF for excision. **a**, Bacterial two hybrid analysis to detect protein-protein interactions. Cells containing fusions of PexA to the T25 subunit of adenylate cyclase CyaA (indicated on left) and Int to the T18 subunit (indicated on top) were spotted on X-gal. Blue colonies indicate a physical interaction between the proteins fused to the CyaA subunits. **b**, Purified 6xHisSUMO-Int (1mM) and/or PexA (0.25mM) were incubated for 30 minutes and then incubated with nickel resin. Unbound and eluted fractions were collected and were run on an SDS-PAGE gel. **c**, Cartoon of *in vitro* excision reaction shows dsDNA fragments containing *attL* and *attR* recombining to *attC* and *attP*. **d**, *In vitro* recombination assay shows when purified PexA or Int are incubated alone with *attL* and *attR*(lanes 1 and 2, respectively), no recombination is detected. As the concentration of PexA increases, recombination products *attC* and *attP* are detected. Uncropped gels are presented in Supplementary Fig. 5. **e**, Cartoon of *in vitro* integration reaction depicting the ability of an RDF to inhibit the activity of an LSR when recombining *attC* and *attP* sites in an integration reaction. **f**, *In vitro* inhibition of recombination (right) by PexA can be seen by the loss of the *attL* and *attR* recombination products as purified PexA increases in concentration.

Recombination in characterized LSR/RDF systems requires solely the LSR, RDF, and DNA substrates^13,17,14^. To determine if PexA directs Int to excise and circularize PLE, *in vitro* recombination assays were performed using PCR fragments containing the *attR* and *attL* sites and purified PexA and Int (Fig. 3c). Addition of neither PexA nor Int alone led to recombination between *attR* and *attL*; however, when Int and PexA were both added, recombination products were detected (Fig. 3c). RDFs have also been shown to block LSR-mediated integration^14^. Consistent with this model, we observed that addition of PexA to an *attC* and *attP* recombination reaction blocks Int-mediated integration *in vitro* (Fig. 3d). These data demonstrate that phage-encoded PexA is the RDF for PLE 1 Int and provide the first example of an LSR/RDF pair being encoded in different genomes.

For PLE to successfully exploit ICP1 PexA for excision, we hypothesized that PexA would be evolutionarily conserved over time. In support of this assertion, analysis of ICP1 genomes from a 12-year period shows that PexA is present and that it is 99% identical in all ICP1 isolates (Supplemental Fig. 4a). The conservation of PexA led us to speculate that although PexA is not essential (Supplemental Fig. 3), it is likely integral to the ICP1 lifecycle. To test this hypothesis, *ΔpexA* phage were competed against wild-type phage in the absence of PLE (Fig. 4a). Strikingly, the proportion of *ΔpexA* phage after each round of passaging dropped significantly, demonstrating that *pexA* is critical for the fitness of ICP1 (Fig. 4b). Hence, ICP1’s ability to avoid stimulating PLE 1 circularization and horizontal spread by acquiring mutations in *pexA* is limited by associated fitness costs. Interestingly, ICP1 *ΔpexA* is still unable to form plaques on a PLE 1 host (Fig. 4c), indicating that PLE circularization is not required for PLE-mediated anti-phage activity.

**Fig. 4.**
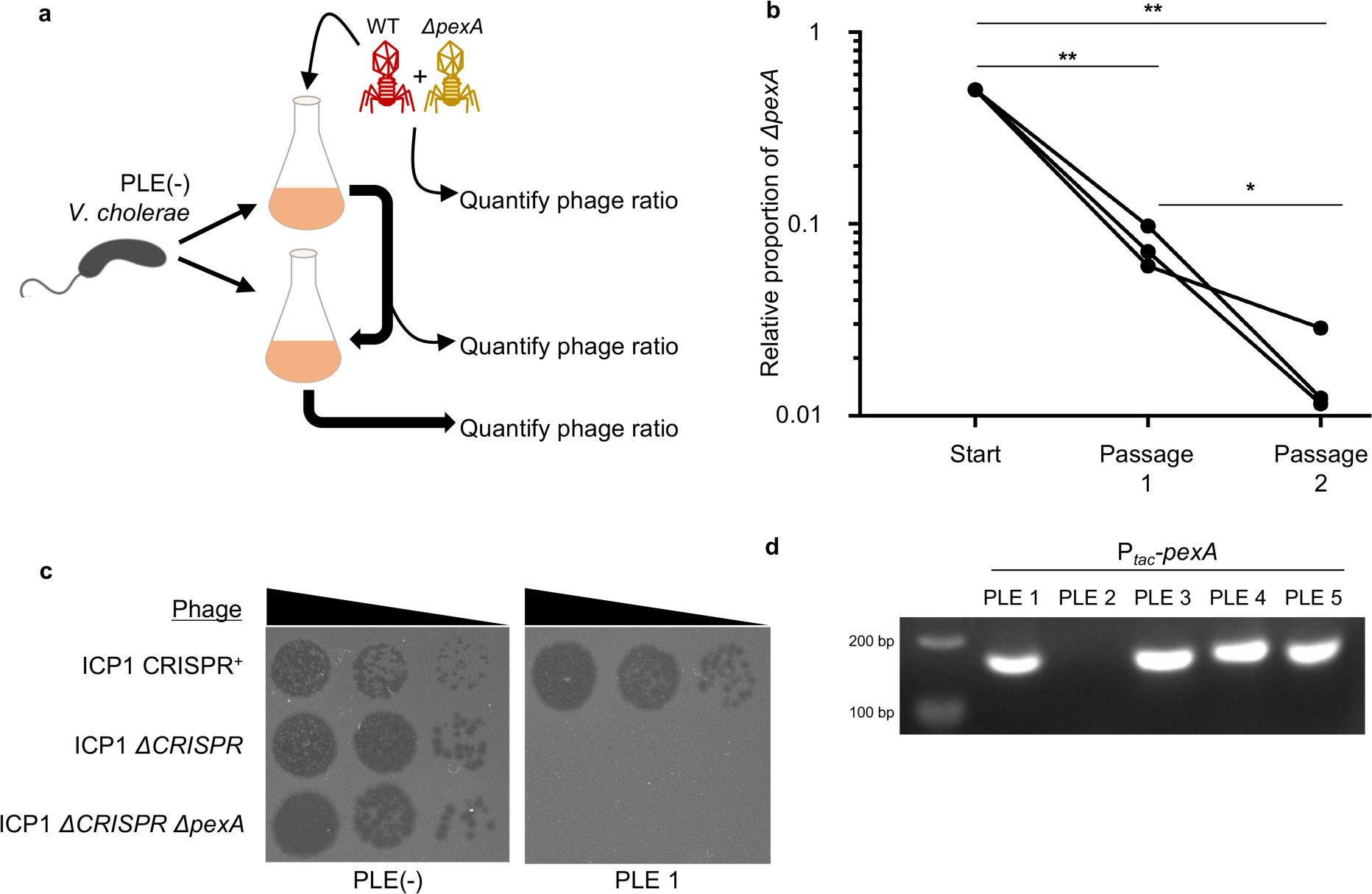
*V. cholerae* PLE Int and ICP1 PexA are in an evolutionary arms race. **a**, Cartoon of fitness competition between ICP1 and ICP1 *ΔpexA*. Mixed phage populations were added to PLE(-) *V. cholerae* at low MOI (0.01) and allowed to infect the culture until lysis. The amount of ICP1 *ΔpexA* in the population was determined, and the lysate was used to infect a new PLE(-) *V. cholerae* culture at low MOI (0.001-0.01). **b**, Three independent mixtures of wild-type ICP1 and *ΔpexA* were passaged, and the relative proportion of *ΔpexA* in the total phage population was calculated at the end of each passage. Each independent mixture was tracked throughout the passage series, and values were normalized to a starting ratio of 0.5 (ratio between 0.3 and 0.6). Significance was calculated from the proportion of *ΔpexA* at each sample point using a Student’s T-Test (*p<0.05; **p<0.005). **c**, Tenfold dilutions of ICP1 spotted on *V. cholerae* PLE+/-lawns showing the ability of different phage strains to form plaques (dark spots, zones of killing). ICP1 and ICP1 *ΔpexA* are unable to form plaques on PLE 1, while the CRISPR proficient phage is able to overcome PLE 1 and form plaques^7^. **d**, Circularization PCR of all PLEs probed during ectopic expression of PexA. The uncropped gel is presented in Supplementary Fig. 5.

As all five PLEs contain a putative LSR and respond to ICP1 infection by circularizing following infection^9^, we set out to examine the conservation of the Int/PexA interaction. Upon ectopic expression of PexA, all PLEs, except PLE 2, demonstrated functional circularization (Fig. 4d). This finding is rather remarkable, given the spatiotemporal distribution of samples from which PLEs have been isolated. For example, the oldest *V. cholerae* isolate known to harbor PLE 5 is an Egyptian isolate from 1949^9^, yet PLE 5 is mobilized by PexA, highlighting the longstanding conflict between ICP1 and PLEs. PLE 2 has the most diverse Int, sharing 25.7% amino acid identity with PLE 1 Int across 63% of the protein, while Ints in PLE 3, PLE 4, and PLE 5 are more similar to PLE 1 Int (Supplemental Fig. 4b). Consistent with the divergence of PLE 2 Int, PLE 2 also integrates into a unique site in the *V. cholerae* small chromosome^9^, indicating that PLE 2 Int recognizes different *att* sites from the other PLEs. Altogether, this data indicates that PLE 2 evolved to recognize a unique RDF, possibly altering *att* site specificity in the process.

We report the first LSR/RDF system in which the interacting LSR and RDF are encoded in separate genomes. The exploitation of PexA by PLE demonstrates co-evolution with ICP1, as PLE has evolved to use an integral phage protein to direct PLE excision and circularization. By binding PexA, which is expressed early in infection, *V. cholerae* PLE 1 is able to quickly detect and respond to ICP1 infection. PLE excision during ICP1 infection enables mobilization of PLE to neighboring cells, and we have previously shown that transduction is Int dependent^9^. It is interesting to note, then, that PLE exploitation of PexA forces ICP1 to directly contribute to horizontal spread of this anti-ICP1 genomic island. As PexA is unique to ICP1, its function as the RDF for PLE 1 Int defines one component underlying the molecular specificity of the interaction between PLE and ICP1. This specificity supports the hypothesis that PLE is not a general phage defense mechanism, but an evolved, highly attuned, and specific response evolved to combat continued predation by ICP1.

Since PLE’s capacity to block ICP1 plaque formation is not dependent on PexA-mediated circularization of PLE during infection (Fig. 4c), we hypothesize that multiple ICP1-encoded signals induce PLE activity, which remain to be elucidated. This model is divergent from the characterized <u>p</u>hage inducible <u>c</u>hromosomal islands (PICIs) in Gram-positive bacteria, in which a single phage protein serves as the only input necessary to de-represses a PICI master repressor, which induces the PICI-encoded RDF and sets off the excision, replication, and packaging cycle^18–20^. Accordingly, phages that avoid inducing PICI activity can be selected for *in vitro*^21^. In contrast, our findings indicate that single evolutionary events that compromise PLE inducing genes in ICP1 cannot prevent PLE induction entirely and helps to explain the evolutionary pressures that lead to the apparent fixation of an active anti-PLE CRISPR-Cas system in ICP1 isolates^7,8^.

## Methods

### General growth conditions

Strains, plasmids, and phage used in this study are listed in Supplementary Table 1-4. Strains were grown with aeration in LB (lysogeny broth) or on LB agar plates at 37°C, unless otherwise noted. Where necessary, cultures were supplemented with streptomycin (100 µg/mL), spectinomycin (100 µg/mL), kanamycin (*V. cholerae* 75 µg/mL, *E. coli* 50 µg/mL), ampicillin, (*V. cholerae* 50 µg/mL, *E. coli* 100 µg/mL), chloramphenicol (*V. cholerae* 2.5 µg/mL, *E. coli* 25 µg/mL), or 5-bromo-4-chloro-3-indolyl-beta-D-galacto-pyranoside (X-gal), 40 µg/mL. Ectopic expression constructs in *V. cholerae* were induced with 1mM IPTG and 1mM theophylline. Phage were propagated on *V. cholerae* hosts using the soft agar overlay method^9^. High titer phage stocks were collected by polyethylene glycerol precipitation^22^.

### Strain construction

PCR products to make chromosomal *V. cholerae* mutants, including chromosomal expression constructs, were created by SOE (splicing by overlap extension) PCR and introduced by natural transformation^23^. Primer sequences are available upon request. Ectopic gene expression vectors were created from a modified pMMB67EH vector^24^ engineered to contain a theophylline inducible riboswitch^25^. Plasmids were constructed using Gibson assembly (NEB), and introduced into *V. cholerae* through conjugation with *E. coli* S17. ICP1 mutants were created as described through CRISPR-Cas gene editing^26^. All constructs were confirmed with DNA sequencing of the region of interest.

### Circularization and excision PCRs

Stool specimens were collected and stored from previous studies^7^. Total DNA was extracted from 100µl stool samples using the DNeasy blood and tissue kit (Qiagen), and 2 µL of extracted DNA was used as template to detect PLE, ICP1, and circularized PLE by PCR. For detection of PLE circularization during phage infection, *V. cholerae* strains were grown to OD_600_=0.3, infected with phage at a multiplicity of infection (MOI) of 5, and allowed to incubate at 37°C with aeration. Samples were taken 5 minutes post infection, boiled for 10 minutes, and 2µL was used as template for PCR using primers depicted in Fig 1a. Resulting reactions were run on a 2% agarose gel and imaged with Gel Green. For Int complementation, ectopic Int was induced for 20 minutes prior to phage infection. To detect PLE excision, 6ng gDNA from uninfected *V. cholerae* and infected *V. cholerae* harvested 15 minutes post infection were used as templates for PCR with primers located in the *V. cholerae* chromosome flanking PLE 1. To detect PLE circularization following induction of ectopically expressed PexA, PLE^+^ derivatives of *V. cholerae* at an OD_600_=0.3 were induced for 5 minutes and samples were boiled and processed for PCR as described above.

### In vivo recombination assay

Constitutively expressed *lacZ* from *E. coli* was engineered such that *lacZ* was flanked by 300 bp containing *att*_*P*_ from circularized PLE and 70 bp containing *att*_*C*_ from a *V. cholerae* repeat (VCR). This construct was integrated into the *V. cholerae* genome at a fixed position (VC2338) by natural transformation. A plasmid with Int or the empty vector control was mated into the reporter strain and individual colonies were picked into 1 mL LB and 2 µL was spotted onto indicator plates with X-gal, IPTG and theophylline.

### PLE circularization screen with mutant phage

Mutant phages, created by targeting the ICP1 genome with CRISPR and collecting viable escape phage^26^, were used to infect the miniPLE *V. cholerae* host using the soft-agar overlay method. Plaques were picked into 50 µL water and boiled for 10 minutes, and 2µL of the boiled template was used for circularization PCR as described above. Since mutants were not clean deletions of individual gene products, hits that failed to circularize miniPLE were verified by making clean deletions of the individual gene products and re-tested for miniPLE circularization phenotypes.

### Western Blot

To detect the presence of PexA following phage infection, *V. cholerae* cultures were grown to OD_600_ = 0.3 and infected with the indicated phage at MOI = 5, and 2 mL samples were taken at the time points indicated. Samples were washed with methanol and phosphate buffered saline (PBS) before being pelleted, re-suspended in 1x Leammli buffer (Bio-rad), and boiled for 10 minutes. Boiled samples were then loaded onto an SDS-PAGE gel for blotting analysis with custom rabbit-α-PexA primary antibodies (GenScript) and goat-α-rabbit-HRP conjugated secondary (Bio-rad). Blots were developed with Clarity Western ECL Substrate (Bio-rad) and imaged on a Chemidoc XRS Imaging System (Bio-rad).

### Bacterial adenylate two-hybrid (BACTH) assay

Genes of interest were cloned into the multiple cloning sites of the pUT18 and pKT25 as described previously^27^. Three independent colonies were separately picked into 1 mL of LB and 3 µL was spotted onto selective medium containing kanamycin and ampicillin with 0.5 mM IPTG and X-gal. Plates were incubated for 24 hours at 30°C before being imaged.

### Protein preparation

*E. coli* BL21 cells containing a pE-SUMO fusion to the construct of interest were grown in LB with antibiotics at 37°C to OD_600_~0.9 and induced for 3 hours with 0.5mM IPTG. Cells were centrifuged and resuspended in lysis buffer (50mM HEPES pH 7.2, 300mM NaCl, 20mM imidazole, Pierce^TM^ Protease Inhibitor Mini Tablets (Thermo), 1mM TCEP, 0.5%Tx-100) and sonicated. Cell debris was removed by centrifugation (18,000x g for 40 minutes at 4°C), and the lysate was applied to a Nickel resin affinity column (HisPur Ni-NTA Resin). The column was washed with two column-volumes wash buffer (50mM HEPES pH 7.2, 1M NaCl, 20mM Imidizole, 1mM TCEP) and eluted with elution buffer (50mM HEPES pH 7.2, 300mM NaCl, 300mM Imidizole, 1mM TCEP). Eluted 6xHisSumo-Int was then run through a HiTrap Heparin HP 5mL column, and pooled fractions were run on a Superose 6 Increase 10/300 GL column on an AKTA Pure 25L system (GE Healthcare). Eluted 6xHisSumo-PexA was run on a Superose 6 Increase 10/300 GL column. To cleave the SUMO tag, 1 µL SUMO protease was added per 100µg of protein and incubated overnight at 4°C. The mixture was then bound to Novex His-Tag Dynabeads and the unbound fraction was collected and analyzed by SDS-PAGE visualized with *Stain*-*Free* technology (Bio-rad).

### In vitro pulldown

Purified 6xHis-SUMO-Int (1000 nM) and/or untagged PexA (250 nM) were added to Novex Dynabeads His-Tag Isolation & Pulldown beads with Binding/Wash buffer (50 mM Sodium-phosphate pH 8, 300 mM NaCl, 0,01% Tween-20) and incubated rocking at room temperature for 10 minutes. Unbound protein was collected and the resin was washed 4x with Binding/Wash buffer. His Elution buffer (300 mM imidazole, 50mM sodium-phosphate pH 8, 300mM NaCl, 0.01% Tween-20) was added, incubated for 5 minutes rocking at room temperature, and collected. Fractions were analyzed by SDS-PAGE visualized with *Stain*-*Free* technology (Bio-rad).

### In vitro recombination

Purified Int and/or PexA were added to 20 µL reactions in the concentrations listed (Fig. 3, Supplemental Fig. 1b) with 200 ng purified PCR products containing the indicated *att* sites in buffer (50 mM HEPES pH 7.2, 150 mM NaCL, 10% glycerol, 0.5 mg/mL BSA, 5mM spermidine). Reactions were incubated for 2 hours at 37°C followed by 10 minutes at 75°C to heat inactivate. The entire 20 µL reaction was then run on a 2% agarose gel and visualized with Gel Green.

### Competition assay

PLE (-) *V. cholerae* was grown to an OD_600_=0.3, and 800 µL was diluted into a total of 5mL LB. Equal PFUs of wild-type and mutant phage were mixed and added at an MOI of 0.01. The OD_600_ of the culture was monitored until lysis. Lysate was collected, treated with chloroform and centrifuged to remove bacterial debris. A second round of passaging was initiated using 10µL of cell free phage-containing lysate diluted 1:100 (MOI=0.01-0.001). The ratio of wild-type to mutant phage in the population was quantified before and after each round of passaging in triplicate. For quantification, two hosts (restrictive and permissive) were used. The restrictive host was CRISPR-Cas^+^ *V. cholerae*^4^ with a spacer targeting *pexA*. On this host, the wild-type phage is unable to form plaques (efficiency of plaquing (EOP) <4x10^−5^), allowing us to enumerate only the *ΔpexA* mutant. The permissive host, without CRISPR targeting, allowed us to quantify the total phage population. The relative proportion of mutant phage was determined to be the amount of mutant phage in the total phage population, and the passaged series was normalized to a starting proportion of 0.5 for visualization (actual values between 0.3 and 0.6). Differences between groups were analyzed using a 2-tailed Student’s T-Test.

### Phage plaque spot plates

*V. cholerae* grown to mid-log was added to 0.7% molten LB agar, poured over a solid agar plate, and allowed to solidify. 3 µL of each ten-fold dilution of phage in LB was spotted onto the solid surface and allowed to dry. Plates were incubated at 37°C overnight before being imaged.

### Genomic analysis

Structure prediction of PLE 1 Int was performed using NCBI Conserved Domain Database^28^ and the MPI bioinformatics toolkit^29^. PexA sequence was displayed with Weblogo^30^. Alignments for PLE Int were performed with PRALINE^31^, and EMBOSS Needle was used to compare PLE 1 and PLE 2 Int^32^.

## Acknowledgements

We would like to thank members of the Seed Lab for thoughtful discussions and feedback. Thanks to Dr. Lindsay Matthews and Dr. Lyle Simmons for the BACTH materials and bacterial strains and plasmids used for protein purification. Thanks to The Glaunsinger Lab for use of their AKTA and specifically Matt Gardner for training on the system. This work was supported by a Kathleen L. Miller Fellowship to ACM from the Henry Wheeler Center for Emerging and Neglected Diseases and grant R01AI127652 to KDS from the National Institute of Allergy and Infectious Diseases (US). KDS is a Chan Zuckerberg Biohub investigator.

## Author contributions

ACM and KDS conceptualized the project, designed experiments and wrote the manuscript. ACM performed the experiments and analyzed the results. KDS supervised the study.

## Competing interests

The authors have no competing financial interests to declare.

## Data availability

Data is presented, and raw competition values available upon request.

